# *De novo* serine biosynthesis is protective in mitochondrial disease

**DOI:** 10.1101/2023.03.23.533952

**Authors:** Christopher B Jackson, Anastasiia Marmyleva, Ryan Awadhpersad, Geoffray Monteuuis, Takayuki Mito, Nicola Zamboni, Takashi Tatsuta, Amy E. Vincent, Liya Wang, Thomas Langer, Christopher J Carroll, Anu Suomalainen

## Abstract

Importance of serine as a metabolic regulator is well known in tumors and raising attention also in degenerative diseases. Recent data indicate that *de novo* serine biosynthesis is an integral component of metabolic response to mitochondrial disease, but the roles of the response have remained unknown. Here, we report that glucose-driven *de novo* serine biosynthesis maintains metabolic homeostasis in energetic stress. Pharmacological inhibition of the rate-limiting enzyme, phosphoglycerate dehydrogenase (PHGDH), aggravated mitochondrial muscle disease, suppressed oxidative phosphorylation and mitochondrial translation, altered whole-cell lipid profiles and enhanced mitochondrial integrated stress response (ISR^mt^), *in vivo,* in skeletal muscle and in cultured cells. Our evidence indicates that *de novo* serine biosynthesis is essential to maintain mitochondrial respiration, redox balance, and cellular lipid homeostasis in skeletal muscle with mitochondrial dysfunction. Our evidence implies that interventions activating *de novo* serine synthesis may protect against mitochondrial failure in the skeletal muscle.

**Bullet points:**

- Serine becomes an essential amino acid in mitochondrial translation defects
- Blocking *de novo* serine biosynthesis promotes progression of mitochondrial disease
- *De novo* serine biosynthesis maintains phospholipid homeostasis upon mitochondrial insult
- Serine biosynthesis sustains redox-balance and mitochondrial translation in disease

## Introduction

Mitochondria are organelles that function at the crossroads of both catabolic and anabolic processes of cellular metabolism (Nunnari & Suomalainen, 2012). The central catabolic function of mitochondria is oxidative breakdown of nutrients to produce ATP. This function is preferential when nutrient availability is low. Upon high nutrient availability, cytoplasmic glycolysis is upregulated for ATP production with mitochondria enhancing production of metabolites, amino acids, and cofactors for anabolic growth (Vasan *et al*, 2020). The biosynthetic pathways are highly tissue- and cell-type specific, making tissues differently sensitive to stress and disease-linked challenges. Therefore, it is not surprising that defects of mitochondrial functions cause an exceptionally broad spectrum of human disorders, ranging from neurodegeneration to heart, muscle and endocrine disorders (Gorman *et al*, 2016; Nunnari & Suomalainen, 2012). The molecular details of disease-related metabolic remodelling and the relevance for disease progression are only starting to be revealed.

Mitochondrial myopathy (MM) is a common form of mitochondrial diseases, typically caused by mutations in genes encoding proteins involved in mitochondrial DNA (mtDNA) expression, replication, or translation (Gorman *et al*, 2016). These defects induce complex stress responses (mitochondrial integrated stress response, ISR^mt^) involving remodelling of cellular anabolic metabolism, including folate cycle, methyl cycle, nucleotide and glutathione synthesis as well as production and secreted metabokines FGF21 and GDF15 in mice and cell cultures (Forsström *et al*, 2019; Khan *et al*, 2017; Nikkanen *et al*, 2016; Kühl *et al*, 2017; Bao *et al*, 2016; Ignatenko *et al*, 2020) and are conserved in human patients (Lehtonen *et al*, 2016; Pirinen *et al*, 2020). Recent data indicate that ISR^mt^ is stagewise, first involving transcriptional activation of ATF5, metabokines and mitochondrial folate cycle, followed by a 2^nd^ stage of upregulated *de novo* serine synthesis and mTORC1 activation (Forsström *et al*, 2019; Khan *et al*, 2014; Nikkanen *et al*, 2016). These previous data have indicated that primary mitochondrial defects cause a remarkable cell-autonomous remodelling of cellular metabolism. Furthermore, the ISR^mt^ cell-non-autonomous response, which occurs via secretion of the metabokines GDF15 and FGF21, modifies fat and glucose metabolism systemically (Tyynismaa *et al*, 2010; Forsström *et al*, 2019). However, whether ISR^mt^ promotes or protects from disease progression *in vivo*, and whether its components can be therapeutic targets, remains unclear.

*De novo* serine biosynthesis (dnSB) is an integral upregulated component of ISR^mt^ in both postmitotic and cultured cells (Forsström *et al*, 2019; Nikkanen *et al*, 2016; Bao *et al*, 2016), as well as a key component to promote anabolic growth in cancer cells (Locasale *et al*, 2011; Pacold *et al*, 2016; Mehrmohamadi *et al*, 2014). The regulatory enzyme of dnSB is 3-phosphoglycerate dehydrogenase (PHGDH, EC1.1.1.95), catalysing the first and rate-limiting step of the three-step process that bifurcates glucose flux from glycolysis to dnSB (Possemato *et al*, 2011; Bao *et al*, 2016; Locasale *et al*, 2011; Vandekeere *et al*, 2018). In addition to its function as proteinogenic amino acid in the muscle, serine can donate donate its one carbon (1C) units via folate cycle for nucleotide synthesis, support glutathione (GSH) and heme synthesis and nicotinamide adenine dinucleotide phosphate (NADPH) synthesis, making serine a major contributor to the cellular redox pool and pentose phosphate pathway (Tajan *et al*, 2021; Mattaini *et al*, 2016; Vandekeere *et al*, 2018). Furthermore, serine supports membrane phospholipid synthesis by providing the head group of phosphatidylserines and serves as a substrate to sphingolipid synthesis (Gao *et al*, 2018).

The reason why serine needs to be synthesized as part of a metabolic stress response, and which of its functions are required upon metabolic stress in different cell types are unknown. Here, we studied dnSB in Deletor mice, a mouse model that manifests with similar findings as patients with adult-onset mitochondrial myopathy: slowly progressive mtDNA mutagenesis and respiratory chain deficiency (Tyynismaa *et al*, 2005). In these mice, *de novo* serine biosynthesis (dnSB) induction is a key component of their disease (Tyynismaa *et al*, 2010; Forsström *et al*, 2019). We report here the consequences of inhibited PHGDH activity in Deletor mice and in cultured cells with mitochondrial dysfunction. We find that *de novo* serine biosynthesis is critical for maintenance of cellular metabolic homeostasis and viability in mitochondrial stress and disease.

## Results

### Inhibition of *de novo* serine biosynthesis aggravates mitochondrial pathology

To test the importance of dnSB for mitochondrial disease progression, we treated Deletor mice, which ubiquitously express a homologous dominant patient mutation in mitochondrial helicase twinkle (Twinkle^dup353-365^)(Tyynismaa *et al*, 2005), and wild type (WT) mice systemically with vehicle or NCT-503, an inhibitor of PHGDH, the rate-limiting dnSB enzyme ((Pacold *et al*, 2016), Figure 1A). Thirty days of daily intraperitoneal injections did not have visible effects on mouse well-being or body weight, indicating no apparent systemic toxicity for NCT-503 as also reported before (Fig. S1A, Pacold et al., 2016).

**Figure 1.**
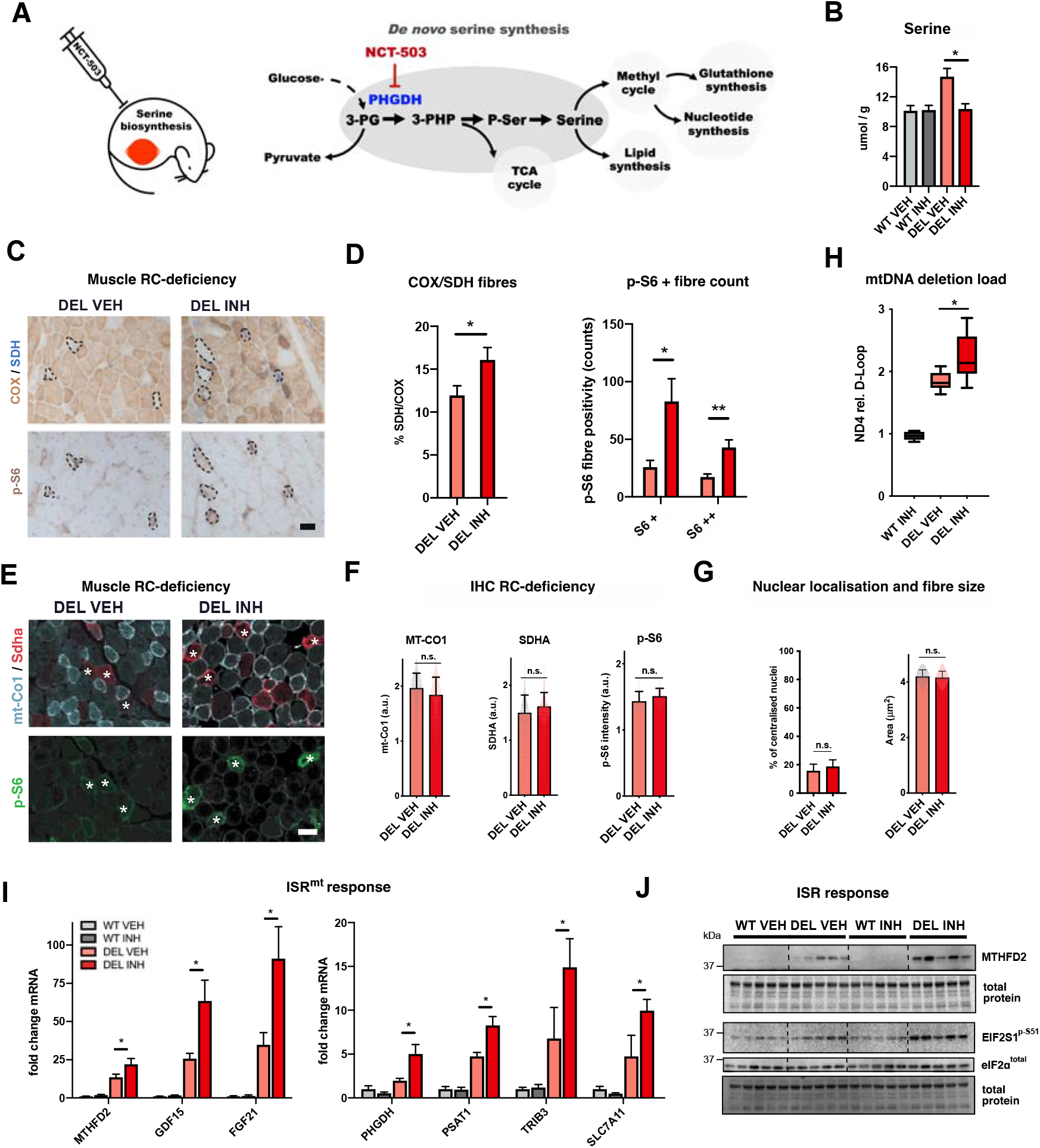
Systemic NCT-503 administration effectively inhibits de novo serine biosynthesis in mitochondrial myopathy. (A) Serine inhibition treatment trial and *de novo* serine synthesis-related cellular pathways. (B) Serine amount upon PHGDH inhibition. NCT-503 administration for 30 days; targeted metabolomics of skeletal muscle (WT VEH n=6, Del VEH n=6, WT INH n=6, Del INH n=8). (C) Effects of PHGDH inhibition for respiratory chain deficiency. Histochemical analysis of combined cytochrome-c-oxidase (COX) and succinate dehydrogenase (SDH) activity. (Brown fibres indicate high COX activity, translucent low COX activity. Blue fibres indicate high SDH activity and mitochondrial proliferation, translucent low SDH activity). Lower panels show immunohistochemical detection of the mTORC1 downstream target phosphorylated ribosomal S6 (S6-P). Images represent sequential frozen sections and black circles matching fibres in subsequent sections. (D) Quantification of RC-deficiency in skeletal muscle from C. Amount of COX negative (COX-) and SDH positive (SDH+) fibres in treated and untreated Deletors (DEL VEH n=6, DEL INH n=7) and amount of strong (+) and very strong (++) p-S6 fibres in treated and untreated Deletors (DEL VEH n=6, DEL INH n=7). (E) Immunofluorescence imaging of Deletor muscle cryosections probed for mitochondrially-encoded protein (Mt-Co1, light blue), mitochondrial mass (Sdha, red) and a myofiber membrane marker (Laminin, white). Lower panels show immunofluorescence for phosphorylated ribosomal S6 (S6-P, green). Scale bar: 100 μm. (F) Quantification of total immunofluorescence of grouped VEH and INH animals from (E). INH show unchanged MT-CO1 levels, SDHA and p-S6 levels. (G) Muscle cross section fibre area quantification and nuclear localisation deduced from (E). All intensities normalised and expressed in arbitrary units (a.u.). (H) MtDNA deletion load; qPCR (ratio of deleted ND4 / intact D-Loop) (WT VEH n=6, Del VEH n=6, WT INH n=6, Del INH n=8). Results normalised to untreated WT mice. (I) Transcriptional induction of ISR^mt^ markers. qPCR; relative to β-actin (WT VEH n=6, Del VEH n=6, WT INH n=6, Del INH n=8). (J) Translational induction of ISR^mt^ marker MTHFD2 and phosphorylation status of eIF2α. Immunoblot (n=5/group). Animals: 25 months old. Bars represent the group average and error bars standard error of mean (SEM), individual animals or skeletal fibres as dots. Statistical significance was performed using ANOVA, * p≤0.05, **p≤0.01. ***p≤0.001. Scale bars 100 µm. Abbreviations: WT=wildtype, DEL=Deletor, VEH=vehicle, INH= NCT-503 inhibitor, PHGDH= D-3- phosphoglycerate dehydrogenase

In the Deletor *quadriceps femoris* muscle, the inhibitor efficiently reduced serine levels to the level of WT mice (Figure 1B), while not affecting the amounts in the WT muscle. These results indicate the efficacy of the drug *in vivo* and the specificity of PHGDH as a regulator of serine levels in mitochondrial disease. Analysis of respiratory chain enzyme activities indicated that PHGDH-inhibition increased the number of respiratory chain (RC)-deficient fibres [cytochrome *c* oxidase deficient (COX negative) and elevated pathology-related succinate dehydrogenase activity (SDH positive)] (Figure 1C,D) in the Deletor muscle. However, COX and SDH protein amounts were not significantly altered (Figure 1E,F). NCT-503 treatment also increased the number of fibres with activated mTORC1 (mechanistic target of rapamycin complex 1; activity readout: phosphorylation of S6 (p-S6), a cytoplasmic ribosome component) (Figure 1D). In wildtype mice we did not observe any COX- or p-S6 activated fibres in treated or untreated animals (Fig. S1B). In Deletors, the total mitochondrial mass in the skeletal muscle and the total fibre size were not significantly affected by the inhibitor treatment, and the proportion of fibers with central nuclei remained the same (Figure 1F,G) but mitochondria appeared distorted (Fig. S1C). Also, multiple mtDNA deletion load increased (Figure 1H). The mtDNA content did not significantly increase in treated Deletors but showed elevated levels compared to WT mice (Fig. S1D). These results show that *de novo* serine synthesis delays generation of mtDNA deletions and progression of mitochondrial pathology.

The transcriptional ISR^mt^ response components (ATF5, MTHFD2, GDF15, FGF21, PSAT1, TRIB3, and SLC7A11) were further induced by the NCT-503 treatment compared to the vehicle (Figure 1I). PHGDH transcription was especially stimulated by the inhibition of PHGDH enzyme activity, suggesting a feedback mechanism (Figure 1I). We tested the effect of NCT-503 by treating an independent Deletor transgenic line (Deletor C-line), which has lower mutant Twinkle expression compared to control mice. The Deletor C-line showed a similar induction for PHGDH upon dnSB inhibition (Fig S1E). The induction of ISR^mt^ was also confirmed on protein level (MTHFD2, Figure 1J). Eukaryotic translation initiation factor 2 subunit 1 (EIF2S1), a key component of the cytoplasmic integrated stress response, showed increased phosphorylation levels (Figure 1J, S1F) and the UPR^mt^ marker HSP70 was significantly increased at the protein level (Fig. S1G). WT mice showed no changes in ISR^mt^ upon treatment, highlighting the specific role of dnSB in the context of disease (Figure 1I). Transcript analysis of muscle atrophy markers, MURF1 and ATROGIN1, showed slightly increased expression after PHGDH inhibitor treatment in Deletor and WT mice (Fig. S1H). However, the muscles expressed no histological signs of cell death.

Altogether, these findings demonstrate vulnerability of muscles with mitochondrial dysfunction to restriction of serine biosynthesis.

### Inhibition of PHGDH alters metabolic profile and lipid balance in mitochondrial myopathy

Next, we analysed the muscle metabolome of healthy and diseased mouse muscle upon dnSB inhibition. Principal component analysis indicated separation of Deletor metabolome from WT, which was further aggravated by NCT-503 (Fig. S2A). For the WT mice, the treatment had little effect (Fig. S2B). NCT-503 significantly affected 1C metabolism (nucleotide and glutathione synthesis, methyl cycle, phospholipid pools) since serine contributes to it in various ways. Nucleotide synthesis intermediates, xanthine and hypoxanthine, accumulated in Deletor mice and these were further increased by NCT-503 (Figure 2A). The highest and most significantly changed metabolite after inhibitor treatment in Deletors was gamma-aminobutyric acid (GABA) shown to accumulate if mitochondrial nucleoside salvage is deficient (Besse *et al*, 2015). The ratio of oxidized to non-oxidized glutathione was increased between Deletors compared to WT mice and non-significantly between treated and non-treated Deletors (Fig. S2C). Apart from serine, the total level of non-essential amino acids was increased in Deletors compared to WT mice but decreased upon treatment (Fig. S2D,E). In the separate analysis of total dNTP pools, dCTP was increased, while the metabolomics indicated decreased steady-state cytosine, potentially as a consequence of altered usage (Fig. S2F). These results, together with the ISR^mt^ induction, supported the conclusion that dnSB is protective in mitochondrial disease *in vivo*.

**Figure 2.**
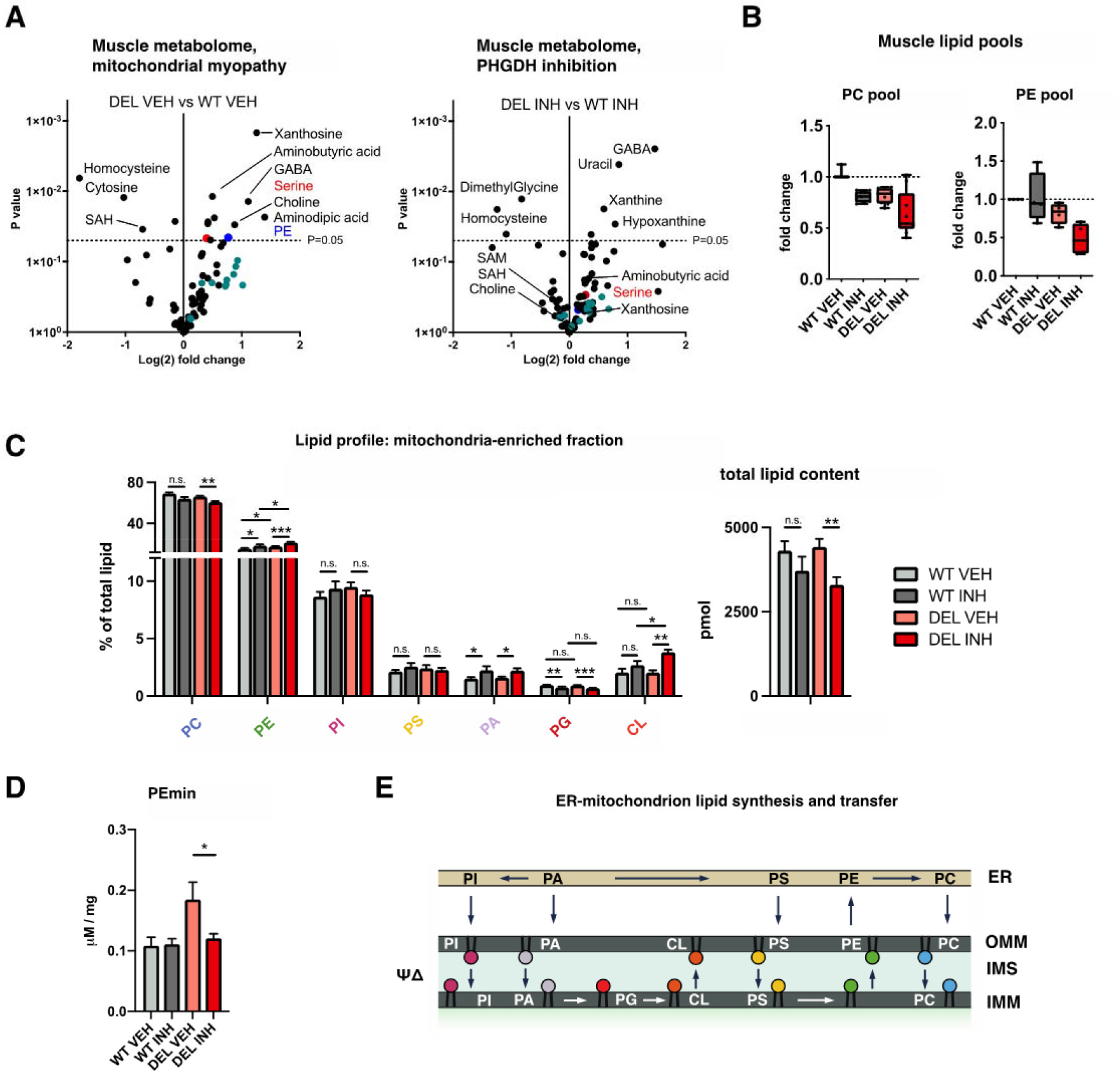
Serine biosynthesis inhibition increases hallmarks of mitochondrial myopathy and alters lipid balance. (A) Metabolomic analysis, skeletal muscle. Volcano plots of treated and untreated Deletors and WT mice. Dots represent individual metabolites. Green:acyl-carnitines. Red: serine. (WT VEH n=6, Del VEH n=6, WT INH n=6, Del INH n=8). (B) Alteration of total PC and PE pools in whole muscle. Untargeted metabolomics. Each dot representing the median value of one fatty acid. (C) Lipid content analysis from mitochondria-enriched fraction from whole muscle (WT VEH n=6, Del VEH n=7, WT INH n=6, Del INH n=8). (D) Phosphoethanolamine (PEmin) level; targeted metabolomics of whole muscle (WT VEH n=5, Del VEH n=7, WT INH n=6, Del INH n=8). (E) Mitochondrial lipid synthesis (adapted from (Mesmin, 2016)). Bars represent the group average and error bars standard error of mean (SEM), individual animals as dots. Statistical significance was assessed using ANOVA, * p≤0.05, **p≤0.01. ***p≤0.001. Abbreviations: phosphatidylcholine (PC), phosphatidylethanolamine (PE), phosphatidylinositol (PI), phosphatidylserine (PS), phosphatidic acid (PA), phosphatidylglycerol (PG) and cardiolipin (CL).

The metabolomic analysis highlighted the importance of *de novo* serine biosynthesis for methylation and phospholipid homeostasis. PHGDH inhibition depleted the Deletor muscle of S-adenosylmethionine (SAM), the major donor of methyl groups to cellular methylation reactions (Figure 2A). The lack of SAH increase (SAM-derived product after methylation event) suggests insufficient synthesis of SAM and overall decreased methylation capacity. This is supported by low dimethylglycine. Intriguingly, in mitochondria, the PE pool was increased (Figure 2C) whereas the total muscle pool was low (Figure 2D). The total and mitochondrial PC pools, however, were depleted by NCT-503 both in healthy and diseased mice (Figure 2B,C). These data emphasize the necessity of dnSB for phospholipid homeostasis in skeletal muscle and especially in mitochondria. Cardiolipin (CL), specific for mitochondrial membranes, increased in amount after PHGDH inhibition in the Deletors, indicating a role for dnSB for CL homeostasis in mitochondria. Next, we asked whether the changed mitochondrial lipid composition affected processing of OPA1, a GTPase controlling mitochondrial membrane dynamics, which binds to CL and is activated in the inner mitochondrial membrane by proteolytic cleavage. However, OPA1 processing was unaffected in whole skeletal muscle homogenates of all our mouse groups (Fig. S2G).

To investigate whether dnSB is a response to potential intra-mitochondrial serine availability, we assessed steady-state level of the mitochondrial serine transporter SFXN1, which were increased in all treatment and non-significantly increased in treated Deletors for SFXN1 (Fig. S2H), suggesting that cell-intrinsic dnSB supports mitochondrial translation upon translation stress.

Together these data show a remarkable remodelling of serine-dependent homeostasis in mitochondrial disease. The evidence indicates that the phospholipid homeostasis is dependent on dnSB especially in mitochondrial disease, as PHGDH inhibition aggravated the findings and elicited a deficiency of the major methyl donor SAM. Interestingly, PE and CL were high in mitochondria while the total cellular pools for PE were low. This may indicate that the lack of SAM hampers PE methylation to PC, thereby accumulating PE to mitochondria (Figure 2E,F). Furthermore, mitochondrial serine uptake and translation appear to be supported by dnSB. The data suggest that dnSB is a protective response in mitochondrial disease progression, fuelling mitochondrial translation capacity and alters the phospholipid homeostasis.

### Serine synthesis is an early response to various mitochondrial stresses

To further understand the relevance of *de novo* serine synthesis in different kinds of mitochondrial insults, we tested temporal treatments of cultured cells with various mitochondrial toxins (Figure 3A): 1) paraquat, causing oxidative stress in mitochondrial matrix; 2) rotenone, complex I inhibitor; 3) oligomycin, F_0_F_1_-ATPase (complex V) inhibitor; 4) antimycin, complex III inhibitor; 5) actinonin and doxycycline, two mitochondrial translation inhibitors; ; 6) FCCP, OXPHOS uncoupler. We also induced mtDNA depletion stress by overexpression of mutants of mitochondrial helicase Twinkle (TWNK) and polymerase gamma (POLG), which lead to a progressive respiratory chain deficiency (Fig. S3A). PHGDH expression (Figure 3B) and increased serine levels (Figure 3C) responded robustly to the different inhibitors of mitochondrial functions. Upstream regulators of ISR^mt^, ATF3-5, were induced after different stressors in cultured myoblasts, ATF3 was especially highly induced upon oxidative stress (paraquat and rotenone), while all ATFs 3-5 responded to actinonin-induced mitochondrial translation stress (Fig. S3B). These results indicate that dnSB induction is a key component of stress responses to different mitochondrial inhibitors and genetic insults of the mtDNA replisome.

**Figure 3.**
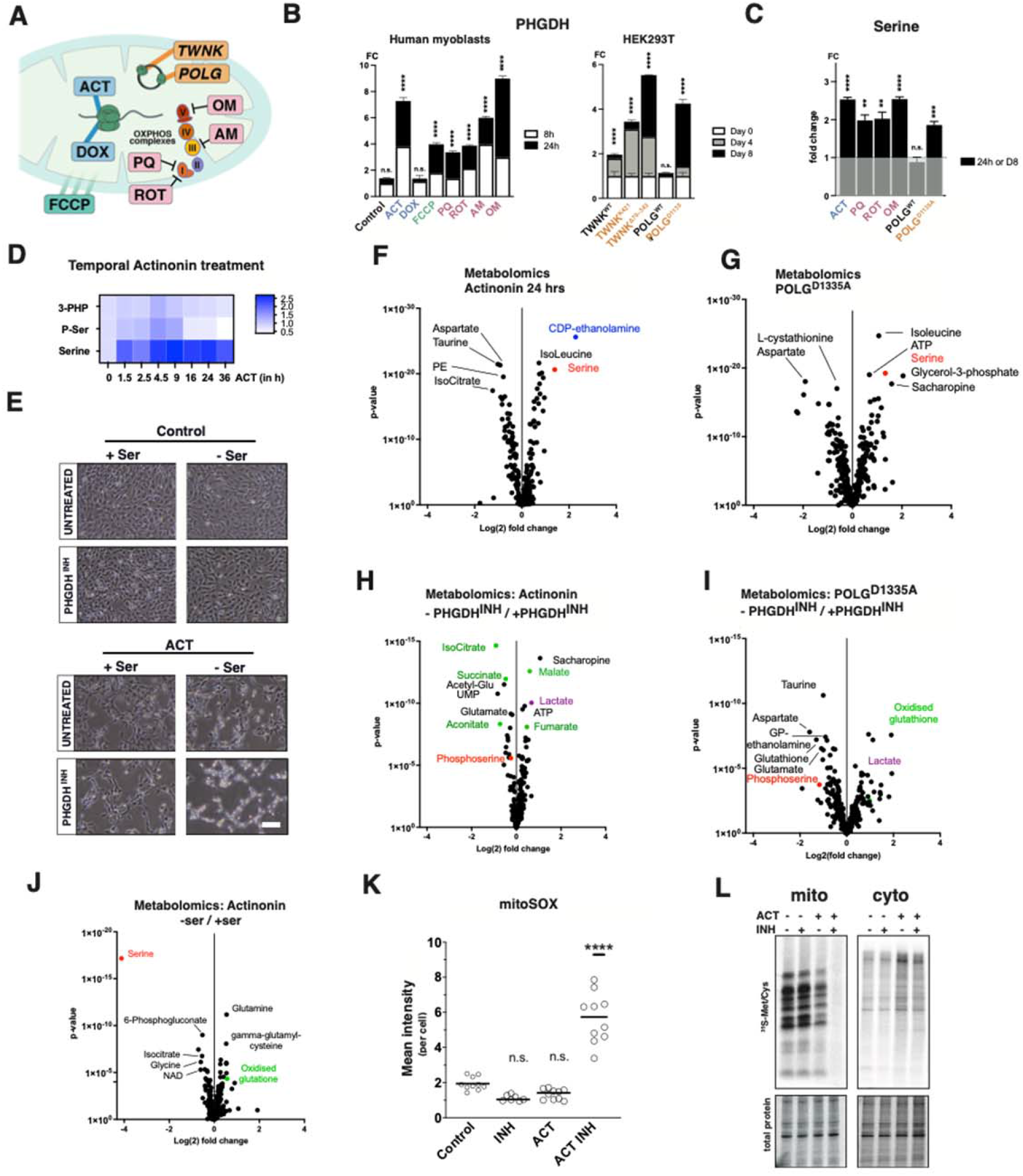
Serine availability and de novo serine synthesis affect cell fitness dependent on mitochondrial dysfunction. (A) Cellular models of mitochondrial dysfunction. Compounds acting on the mitochondrial ribosome (ACT, DOX), protonophore acting on membrane potential in green (FCCP), compounds directly affecting the respiratory chain in red (PQ, ROT, AM, OM), genetic models of dysfunction by mutant overexpression (TWNK, POLG), (B) PHDGH expression transcript levels. Left panel: human myoblasts with indicated treatment for 8 and 24 hrs, right panel: HEK293T cells with overexpressed POLG, WT or mutant TWNK for 4 to 8 days. Normalization against corresponding non-treated or uninduced controls. Error bars represent triplicate measurement, SD. (C) Serine levels in cells treated for either 24 hours for toxins or 8 days for POLG mutants normalized to respective untreated controls. Values determined from untargeted metabolomics. Error bars represent SD of biological quintuplicates. (D) Heatmap of temporal actinonin treatment with indicated timeline of serine *de novo* metabolites 3-phosphohydroxypyruvate (3-PHP), 3-phosphoserine (P-Ser) and serine. Untargeted metabolomics. Intensities calculated from biological quintuplicates. (E) Growth experiments under serine containing and serine deprived conditions in C2C12 myoblasts after 24 hours of drug treatment. Scale bar: 1000 μm. DIC imaging. (F) Actinonin-treated (24 h) C2C12 myoblasts compared to untreated controls. Untargeted metabolomics. (G) POLG^D1135A^ mutant overexpression (4 d) compared to POLG^WT^ expression. Untargeted metabolomics. (H) Untargeted metabolomics; effect of short-term (2.5 h) inhibition of mt-translation with or without NCT-503, compared to actinonin-only treated cells. C2C12 cells. (I) Untargeted metabolomics of POLG^D1135A^ mutant overexpression (4 d) with and without *de novo* serine inhibition. Volcano plot. (J) Untargeted metabolomics. Effect of short-term (2.5 h) inhibition of mt-translation with or without serine-deprivation compared to actinonin-only treated cells. (K) Mitochondrial oxidative stress upon inhibition of mt-translation and/or *de novo* serine biosynthesis. MitoSOX-probe detection. (L) Mitochondrial and cytoplasmic translation products, ^35^S-metabolic pulse labelling. Actinonin and NCT-503 treatments as in E. All metabolic data points represent mean of biological quintuplicates of untargeted metabolomics; mRNA level from biological triplicates. Abbreviations:ACT=actinonin,DOX=doxycycline, PC=phosphatidylcholine, PE=phosphatidylethanolamine, PHGDH=D-3-phosphoglycerate dehydrogenase, PQ=Paraquat, OM=oligomycin, ROT=Rotenone.

We then asked whether dnSB is an early or late component in ISR^mt^ progression in cultured cells (primary diploid human myoblasts) and chose to use actinonin as a stressor for the mtDNA expression system, sharing closely the stress-type also in the Deletor mouse muscle (Richter *et al*, 2013). Untargeted metabolomics analysis of actinonin-treated cells revealed a time-dependent decrease in PSAT1-dependent phosphoserine, while serine was increased only 1.5 hrs after exposure (Figure 3D). Similar to the results from skeletal muscle, the phospholipid pools (PE and PC species) decreased after 16 hrs of exposure, while PE precursor started to accumulate at that time (Fig. S3C,D). The ultrastructure of the inhibitor-treated cells showed complete loss of cristae with appearance of myelinosome-like membranous lipid aggregates (Fig. S3E). Furthermore, short-term exposure to actinonin resulted in a shift to increased phosphorylation of ribosomal subunit S6 (p-S6) levels, a target of mTORC1, with a similar timeline as that of nucleotide metabolism and aspartate increase. Decline of mTORC1 was accompanied by increase of ATF4 activation (Fig. S3F). These results demonstrate that mt-translation stress involves dnSB as an early response.

Serine can be derived from both exogenous and endogenous sources (Locasale et al., 2011; Mullarky et al., 2016). We tested the effect of dnSB upon mitochondrial dysfunction by eliminating the exogenous source, by pre-culturing the cells in serine-free media for 24h. In healthy cells, an additional 24-hour treatment with NCT-503 showed no visible phenotype. However, in cells with mitochondrial translation stress, dnSB inhibition caused cell death. Under PHGDH inhibition, media-supplied serine alone showed only minor compensation for dnSB loss, while D^3^-serine uptake by cells, measured from medium and intracellular serine amount, had a similar uptake capacity in actinonin-treated and untreated cells (Fig. S3G). These results emphasize that cells with mitochondrial translation stress depend on serine availability (Figure 3E). Actinonin incubation only mildly affected cell viability after 24 hrs, when assessed by markers of necrosis (PI) and apoptosis (Annexin V) and was not exacerbated by NCT-503 (Fig. S3H). These data clearly demonstrated that mitochondrial dysfunction causes metabolic adaptation dependent on endogenous serine synthesis and proved *de novo* serine synthesis to be essential for cell survival under mitochondrial stress.

Next, we investigated how the metabolome is affected by serine deprivation. We first triggered short-term mitochondrial translation stress via actinonin (C2C12 cells) or genetic mutation (HEK cells overexpressing a POLG mutant). Both conditions, showed increased serine levels with partially distinct lipid signatures and aspartate depletion (Figure 3F,G). When dnSB was inhibited in these mt-stress conditions, phosphoserine decreased, and TCA cycle intermediates were altered (Figure 3H,I). The PE and PC lipid profile showed a decrease with dnSB inhibition alone, exacerbated at 6h and partial recovery at 24h with additional mt-translation stress (Fig. S3I). Lactate increase indicated a shift towards glycolytic and glutamine-dependent metabolism, accompanied by a strong increase in oxidised glutathione already at 2.5 hours of serine withdrawal (Figure 3J). The intermediates of the transsulfuration pathway decreased, with consequent decrease of glutathione, impaired redox balance and oxidative stress. Indeed, mitochondrial ROS, assessed by mitoSOX signal, remarkably increased in cells with inhibition of both mt-translation and dnSB, but not in either treatment alone (Figure 3K, S4A). In addition, NCT-503 worsened the actinonin-induced mitochondrial respiration defect, even if serine was supplemented in the media (Fig. S4B). This was also reflected in the loss of mitochondrial membrane potential (Fig. S4C,D). To further test the on-target effect of NCT-503, we generated a PHGDH knockout in HEK293 cells (PHGDH^KO^, (Fig. S4E). NCT-503 inhibited growth of WT and PHGDH^KO^ HEK293 cells, as reported before (Pacold et al., 2016). If further challenged by mitochondrial translation inhibition by actinonin, the PHGDH^KO^ cells were not viable in actinonin-treated conditions (Fig. S4F). These results indicate that newly / on-site synthesized serine is required for maintenance of mitochondrial translation upon metabolic stress.

These results prompted us to assess translation capacity by ^35^S-pulse labelling of *de novo* mitochondrial protein synthesis. The cells were pre-treated for 24 hours (actinonin plus/minus; serine plus/minus) prior to the assay but were left in drug-free media for the length of labelling. Actinonin expectedly decreased mitochondrial translation rates with a rapid recovery when the drug was withdrawn. Strikingly, however, inhibition of *de novo* serine synthesis completely blocked the cellular ability to reinitiate mitochondrial protein synthesis (Figure 3L). Cytoplasmic translation capacity was unaffected by these treatments (Figure 3L). These data indicate that dnSB provides serine especially to mitochondria in stress, to maintain mitochondrial translation.

In summary, these data demonstrate that the non-essential amino acid, serine, becomes essential under variety of conditions of mitochondrial stress. It maintains basic cellular functions including lipid synthesis and specifically supports translation of mtDNA-encoded proteins and keeps up oxidative functions.

## Discussion

Here, we report that serine becomes an essential amino acid in cells and tissues with mitochondrial translation stress. Our data show that intracellular *de novo* serine biosynthesis has a critical role in maintaining homeostasis in muscle with mitochondrial disease. In cultured cells with different kinds of mitochondrial dysfunctions, dnSB is a first-line responder to mitochondrial stress. Remarkably, dnSB supports mitochondrial respiration and maintenance of redox-state, nucleotide homeostasis, and phospholipid pools. Inhibition of dnSB in mitochondrial myopathy mice aggravates histopathological and molecular hallmarks in the skeletal muscle. Importantly, in mitochondrial respiratory chain deficiency, exogenous serine, which can be obtained from food or breakdown of proteins, is not able to compensate the lack of dnSB. Our data indicate that local supply of newly synthesized serine is essential to maintain mitochondrial translation and oxidative functions in tissues and cells with mitochondrial dysfunction.

The roles of serine as regulator of stress responses are becoming increasingly clear (Bao et al., 2016; Forsström et al., 2019; Khan et al., 2017; Kühl et al., 2017; Nikkanen et al., 2016). Serine is an important one-carbon (1C) donor for folate-driven 1C-cycle, the major anabolic pathway for cell growth and repair. The 1C cycle drives e.g., glutathione, nucleotide, and phospholipid synthesis, and provides methyl-units for epigenetic DNA or histone modifications. Its roles are crucial for rapidly growing cancer cells (Pacold *et al*, 2016; Locasale *et al*, 2011; Maddocks *et al*, 2017; Ngo *et al*, 2020; Muthusamy *et al*, 2020; Reid *et al*, 2018; Samanta *et al*, 2016). Our data indicate that both postmitotic and proliferating cells with mitochondrial translation stress use similar, cancer-cell-like anabolic pathways for restoration of cellular homeostasis. In a disease situation, serine drives essential repair and stress response pathways, including glutathione synthesis, phospholipid synthesis and maintenance of nucleotide homeostasis. Serine has also been suggested to be the limiting amino acid for mitochondrial translation (Sullivan *et al*, 2019; Tajan *et al*, 2021). An essential role of dnSB may indeed be to secure serine supply for the disease-crippled mitochondrial translation, probably both via serine-dependent formylation of the initiator methionine and by direct L-serine supply for translation (Kory et al., 2018). Our results indicate that dysfunctional mitochondria become dependent for newly synthesized serine.

NCT-503 has not been found to cause major toxicity for healthy mice (Pacold et al., 2016; Padrón-Barthe et al., 2018). This is surprising, as serine – in contrast to other amino acids - is directly used for synthesis of phosphatidylserine and ceramide species (hydrophobic moiety of sphingolipids) that are important mediators of signalling cascades involved in apoptosis, proliferation, and stress responses (Gao *et al*, 2018; Kay & Fairn, 2019). Blocking of dnSB or extracellular serine supply was reported to induce production of cytotoxic deoxy-sphingolipids and reduced ceramide pools but was not found to have an effect on total PC or PE pools in cultured cells (Gao *et al*, 2018). Our data indicate, however, that NCT-503 causes quite remarkable changes in phospholipid pools even in healthy aged mice, further aggravated by mitochondrial dysfunction.

In conclusion, our evidence highlights the importance of serine-directed pathways in maintenance of mitochondrial translation, oxidative functions, redox balance, and phospholipid synthesis in both growing cells and in mitochondrial disease. Inhibition of *de novo* serine synthesis in mitochondrial dysfunction changes fuel utilisation and modifies anabolic outputs of lipid synthesis and redox capacity in response to mitochondrial dysfunction. Our data indicate that mere serine supplementation cannot rescue the mitochondrial need for this amino acid in disease, but approaches boosting *in situ* serine synthesis in mitochondrial disease could attenuate the disease progression. These findings propose that the intracellular localization of serine biosynthesis is critical for survival in metabolic stress.

## Star Methods / Experimental procedures

Further information and requests for resources and reagents should be directed to and will be fulfilled by the Lead Contacts.

### Ethical approval

The National Animal Experiment Review Board and Regional State Administrative Agency for Southern Finland approved the animal experimentation, which were conducted according to the European Union Directive. Maintenance of mice was performed under licences: ESAVI/11682/04.10.07/0217.

### Animal Models and PHGDH inhibition

The Deletor mice express a dominant in-frame duplication, homologous to a human progressive external ophthalmoplegia (PEO) mutation. Deletor mice were used as an *in vivo* model system of mitochondrial dysfunction. Deletors ubiquitously overexpress mitochondrial Twinkle helicase carrying a human patient mutation. Due to the mtDNA replication defect, these animals accumulate mtDNA deletions which lead to late onset muscle myopathy (Tyynismaa *et al*, 2005). Two previously reported transgene lines were used in this study (Tyynismaa *et al*, 2005). The lines differ by severity of the disease: D line showing a stronger myopathy phenotype as compared to the C line. Age- and sex-matching wild type animals were used as controls. Daily injections of the vehicle and the compound were performed intraperitoneally to 24-months-old male mice. Treatment length of 30 days. NCT-503 was freshly prepared prior to the injections. 20 μM stock was dissolved in EtOH (5%), PEG400 (35%) and 30% Cyclodextrin (60%) and was administered to the mice at serum-stable concentrations of 40 mg/kg (Pacold *et al*, 2016).

### Cell culture and manipulations

Mammalian cells with mitochondrial RC dysfunction are metabolically dependent on glycolysis to source ATP (Robinson *et al*, 1992), on pyruvate for aspartate synthesis (Birsoy *et al*, 2015) and pyrimidines (King & Attardi, 1989) and uridine (Löffler *et al*, 1997), therefore all experimental conditions contained glucose, pyruvate and uridine as indicated.)

Murine C2C12 myoblast cells were grown in MEM (Gibco, 11095) with or without L-serine (Sigma, S4311; 42.0 mg/L). Culturing of primary diploid human myoblasts was performed as described before (Khan *et al*, 2017). If not indicated otherwise, cells were treated in 150 μM actinonin for indicated time points ±NCT-503 and ±Serine. Inhibitor treatments were done for indicated time points: Paraquat (Sigma, 36541), oligomycin (Sigma, O4876) 1 μl stock Oro for 24h or indicated time points, FCCP (Sigma, C2920) 1 μl for 15 minutes or indicated, antimycin A (Sigma, A8674) 1μl or indicated time point, rotenone (Sigma, R8875) 1 μl or indicated time point were applied.

CRISPR-mediated PHGDH knockout was performed in HEK cells using the protocol previously described (Hilander et al., 2022, Biorxiv preprint v2) using two single guide (sg)RNA approach targeting Exon 1 of PHGDH (1: 5’-ACTCACAGCGGCCGATTCCG-3’ and 5’-ACTCACAGCGGCCGATTCCG-3’).

Growth curves for cumulative differences in growth under actinonin treatment was performed over 72 hours and values plotted das the cumulative population doubling as follows:

Population doubling = (log(cell number Day 3) - log(cell number Day 0))/log(2)

### Cytoplasmic and mitochondrial translation labelling

Mitochondrial translation assays were performed with EasyTag Express ^35^S protein labeling mix (PerkinElmer, #NEG772014MC). Assays were performed in cells grown on 60 mm dishes, pretreated with drugs for 24 h. Cells were incubated for 30 minutes in 2 ml of pre-warmed media consisting of DMEM without methionine or cysteine (Sigma #D0422) supplemented with 1x glutamax, 10% dialyzed FBS and 1 mM sodium pyruvate. To inhibit cytoplasmic translation anisomycin (Sigma #A9789) was added to the cells to a final concentration of 100 μg/ml five minutes prior to the addition of the EasyTag labelling mix. For cytoplasmic labelling, chloramphenicol (40 μg/ml) was added 30 minutes prior to labelling. For pulse labelling cells were washed, incubated with the labelling mix and harvested after 30 minutes. Protein extracts were separated on 12% stain-free polyacrylamide gels (Bio-Rad), the gels were dried and exposed to a phosphor imaging screen. Phosphor imaging screens were imaged using a FLA-7000 Typhoon scanner (GE Healthcare) and total protein was imaged using a Chemidoc imaging system (Bio-Rad).

### Mitochondrial isolation

Mitochondria were isolated from quadriceps femoris (QF) from Deletors to enrich for mitochondrial protein. Briefly, QF was homogenised in a Potter-Elvehjem with a rotor-driven pestle in HIM buffer (10 mM HEPES-KoH, pH7.6, 100 mM KCl, 3 mM MgCl2, 0.1 mM EDTA, 10% glycerol) supplemented with 1 mM PMSF and protease inhibitors (Sigma). Briefly, one half of the QF was lysed in 5 ml HIM buffer, strained with a 100 μm-pore cell strainer and the sieve rewashed with 5 ml homogenisation buffer. Mitochondrial fractions were obtained by differential centrifugation at a low step at 700xg for 20 minutes and resulting supernatant at 10’000xg and was repeated. All steps were performed at 4°C.

### DNA manipulations and quantitation

Total genomic DNA was extracted by proteinase K digestion following standard phenol–chloroform extraction with ethanol precipitation or column-based extraction (Macherey-Nagel). MtDNA deletion load was assessed with a modified triplex assay by measuring D-Loop, ND1 and ND4 simultaneously (Khan *et al*, 2017; Rygiel *et al*, 2015). MtDNA copy number was determined by real-time quantitative PCR (qPCR) as described before (human: Jackson et al., 2012; mouse: Kahn et al., 2017) with primers indicated in Table S1. All RT-qPCR measurements were performed on a CFX96 Touch qPCR system (Biorad) using IQ SybrGreen kit (Biorad) as previously described (Forsström *et al*, 2019). Primers sequences are available (Table S1).

### Gene expression

RNA was extracted by standard TRIzol (Invitrogen) chloroform precipitation from snap-frozen tissues homogenized by a Fast-Prep w-24 Lysing Matrix D (MP Biomedical) and Precellys w-24 (Bertin Technologies) apparatus. Total RNA from cells was extracted using a miRNA kit (Qiagen) according to the manufacturer’s instructions. DNase digestion was either in eluted RNA or on-column. Usually, 2 μg of total RNA was reverse transcribed using the Maxima first strand cDNA synthesis kit (ThermoFisher). Quantitative real-time PCR amplification of cDNA was performed with IQ SybrGreen kit (Biorad) on CFX96 Touch qPCR system (Biorad). Relative expressions were normalised to β-actin or 18S rRNA. The linear range and specificity of all primer pairs was determined by serial dilutions and sequencing.

### Immunoblotting

Total protein was extracted with RIPA buffer using phosphatase inhibitors tablets (ThermoFisher Scientific) and sodium orthovanadate to preserve phosphorylation sites. Protein extracts from tissue were prepared as described (Forsström *et al*, 2019). Briefly, tissues were homogenised in 1xPBS containing Triton-X and dodecyl maltoside (f.c.1%) in a Precellys w-24 bead homogenizer (Bertin Technologies), incubated on ice for 30 minutes and collected by centrifugation at 20’000xg for 20 minutes. Protein concentrations were determined using Bradford assay. Proteins were separated on appropriate gels by SDS-PAGE usually on 4-20% gradient gels (Biorad), transferred to PVDF membranes and blocked with 5% milk in 1xTBS-T for 1 hour and primary antibodies incubated overnight at 4°C in1:1000 in 3% BSA in TBS-T buffer. Antibodies used were: MTHFD2 (Abcam, #ab37840), SDHA (Abcam, ab14715), p-S6 (Cell signalling, #4858), total S6 (Cell signaling, #2217), and β-ACTIN (Santa Cruz, #1616), TOM20 (Santa Cruz, #11415), PORIN/VDAC1 (Abcam, #ab15895), HSP70 (Abcam, #ab53098), α-TUBULIN (Abcam, #28439), EiF2α (Invitrogen, #4478G), PHGDH (Proteintech, #14719-1-AP), EIF2S1 (p-S51) ab32157, SFXN1 (Sigma, # HPA019543).

### Microscopic structural analyses

Mitochondrial staining was performed as described before (Forsström *et al*, 2019). Briefly, the cells were stained for 15 minutes in 50 nM Mitotracker CMXRos, rinsed with 1xPBS and fixed in ice-cold acetone and mounted in VectaShield. Immunohistochemistry and immunofluorescence staining were performed as described before (Forsström *et al*, 2019). Samples were imaged in a Zeiss Observer 2.1 equipped with ApoTome technology. Image contrast was adjusted within the software of the microscope. For electron microscopy samples were fixed in 2% GA in phosphate buffer and routinely processed by routine protocols. Contrast enhancement was performed using iTEM software and Photoshop 23.5.1.

### Immunofluorescent labelling of muscle tissue

Mitochondrial immunofluorescent labelling was performed using a protocol adapted from (Rocha *et al*, 2015). Briefly, 10 μm cryosections were thawed at room temperature for 1 hour, before fixing for 3 minutes in 4% PFA. Sections were washed, and then permeabilised using an ascending and descending methanol gradient (70% for 10 minutes, 95% for 10 minutes, 100% for 20 minutes, 95% for 10 minutes and 70% for 10 minutes). Sections were washed with TBST and blocked with 10% NGS for 1h. Following further TBST washes sections were blocked using a Vector Shield Avadin-Biotin blocking kit. Sections were blocked with Avidin D solution for 15 minutes and Biotin solution for 15 minutes, with TBST washes after each. Sections were then blocked with a Vector Shield Mouse on Mouse (MOM) blocking kit following the manufacturer’s instructions. Anti-Mt-Co1 (Abcam ab14705, 1:100), anti-SDHA (Abcam ab14715, 1:100) and anti-Laminin (Sigma-Aldrich L9393, 1:50) were diluted in MOM dilutant and applied to sections before incubation overnight at 4°C. The next day sections were washed with TBST and secondary antibodies Goat Anti-mouse IgG2a Alexa 488 (Life Technologies A21131, 1:200) and Goat anti-Mouse IgG1 biotin (Life Technologies A10519, 1:200) were diluted in MOM dilutant and applied to sections before incubating for 2h at 4°C. Sections were washed with TBST and then Streptavidin Alexa 647 (Life Technologies S32357, 1:100) and DAPI (1:400) were diluted in MOM dilutant and applied to the sections before incubation 2h at 4°C. Sections were washed and mounted in prolong Gold antifade mountant before imaging.

Serial sections to those used for mitochondrial labelling were labelled for p-S6. Briefly, sections were thawed at room temperature for 1 hour, before fixing with methanol free 4% PFA for 15 minutes. Slides were rinsed in PBS and sections blocked with 1xPBS/5% NGS/0.3% Triton for 1 hour at room temperature. Sections were blocked with an Avidin-Biotin blocking kit as described above. Antibodies p-S6 (Cell signalling, #4858, 1:100) were diluted in 1xPBS/1%BSA/0.3%Triton and incubated overnight at 4°C. Following washes, secondary antibody Goat anti-Rabbit Alexa-488 (Life Technologies A11006, 1:200), was diluted in 1xPBS/1%BSA/0.3%Triton, and applied to sections before incubation for 2 hours at 4°C. Sections were washed before mounting in prolong Gold and imaging.

Imaging of both experiments was completed using an AxioImager Z1 with motorised stage and sections were tiled using Zen Blue edition. For the mitochondrial section single muscle fibres were segmented automatically using an inhouse software as described previous (Ahmed *et al*, 2017). This allowed us to export an average intensity for mtco1 and SDHA and muscle fibre area. The number of central nuclei were counted and expressed as a percentage of the total fibres analysed. For the p-S6 section fibres were manually segmented in the same software to export mean p-S6 intensity per fibre. Comparison of serial sections allowed us to assess the levels of pS6 at a single cell level in the context of mitochondrial mt-Co1 and SDHA.

### TMRM and mitoSOX stainings

To measure mitochondrial membrane potential tetramethylrhodamine methyl ester probe (Sigma-Aldrich) was used. Reactive oxygen species were assayed using MitoSox (Thermo Fisher Scientific). Cells were stained 30 minutes with TMRM or 15 minutes for MitoSOX. FCCP was used as positive control. Cells were washed with 1xPBS and imaged in phenol-red free medium using an Evos microscope system. Quantitation was performed using CellProfiler.

### FACS analysis for cell cycle and viability

Cell cycle and apoptosis were assayed using propidium iodide (PI) with subsequent FACS sorting. Apoptosis was assayed using a combined PI and AnnexinV staining following the manufacturer’s instructions. All cells were sampled at 24 hours incubation with the indicated concentrations, if not mentioned otherwise. Samples were run on a BD Science Accuri FACS machine. Subsequent data analysis and image generation was performed using FlowJo.

### Histological staining

Histological stainings for COX/SDH were performed as described in detail in (Nikkanen *et al*, 2016; Forsström *et al*, 2019).

### Targeted metabolomics

Targeted metabolomic analysis was performed as before (Forsström *et al*, 2019). Briefly, 20 mg of tissue was homogenised in a Precellys w-24 (Bertin Technologies) apparatus with added 20 μl of internal labelled standard mix and 500 μl of pure acetonitrile (ACN) and 1% formic acid (FA) with second step extraction of 500 μl of 90/10% ACN/H_2_O+1%FA.

To assess serine flux, cells were cultured in serine-less media with labeled D3-serine ((98%; DLM-582-0.1, Cambridge Isotope Laboratories) or unlabelled L-serine at a final concentration of 285 μM for 6 hours and media and cells collected. Cells were washed once in 1xPBS and metabolites extracted by direct scraping in a precalculated volume of ice-cold methanol/ACN/H_2_O. All samples were analysed and quantified using WATERS XEVO-TQ-S triple quadrupole mass spectrometer coupled to ultra-pressure liquid chromatography.

### Untargeted metabolomics

Untargeted analysis and cold metabolome extraction was applied for metabolomics from cells. Briefly, cells were grown on 6-well dishes and collected by washing twice in 75 mM ammonium citrate buffer (pH 7.4) and immediately flash frozen in liquid nitrogen. Samples were extracted by extraction buffer (ACN:methanol:water) at −20°C and analysed a described before (Fuhrer *et al*, 2011). Muscle tissue samples were subjected to a hot extraction protocol. Briefly, approximately 20 mg of snap-frozen tissue were homogenised in 500 μl 70% (v/v) ethanol in water (extraction solvent) in a pre-cooled Precellys w-24 (Bertin Technologies) apparatus with the use of ceramic beads. Lysed sample was transferred to a 15 ml tube with an additional wash with 500 ul extraction solvent for a total of 1 ml. During these steps, samples were kept at −20°C in an ethanol cold bath at all times. Hot extraction was performed by addition of 75°C warm extraction solvent to the lysed samples and incubated for 1 minute. After extraction, samples were cooled to −20° for storage. For analysis, samples were vacuum-dried, resuspended in water prior to analysis. Extracted samples were analysed on an Agilent 6550 QTOF instrument in negative mode. Putative annotations were created by matching to Human Metabolome Database v.3.6. using mass per charge (0.001 m/z tolerance) and isotopic correlation patterns. Untargeted and targeted data was analysed using MetaboAnalyst 3.0 software in interquatile range, autoscaled, log-transformed and missing values estimated by KNN (Xia *et al*, 2009). Testing between groups was performed using univariate analyses, multiple t-testing and 2-way ANOVA. Separation was tested in unsupervised multivariate, partial least squares discriminant (PLSDA) analysis or principal component analysis (PCA). Metabolites from targeted metabolomics were rated significant if p≤0.05. Metabolites from untargeted metabolomics were rated significant if p≤0.01.

### Quantitative mass spectrometry of glycerophospholipids

Mass-spectrometric analysis was performed essentially as described (Tatsuta, 2017; Özbalci *et al*, 2013). Lipids were extracted from mitochondria-enriched fraction in the presence of internal standards of major phospholipids (PC 17:0-20:4, PE 17:0-20:4, PI 17:0-20:4, PS 17:0-20:4, PG 17:0-20:4, PA 17:0-20:4, all from Avanti Polar Lipids) and CL (CL mix I, Avanti Polar Lipids). Extraction was performed according to Bligh and Dyer with modifications and analyzed on a QTRAP 6500 triple quadrupole mass spectrometer (SCIEX) equipped with nano-infusion splay device (TriVersa NanoMate with ESI-Chip type A, Advion). Mass spectra were processed by the LipidView Software Version 1.2 (SCIEX) for identification and quantification of lipids. Lipid amounts (pmol) were corrected for response differences between internal standards and endogenous lipids. Correction of isotopic overlap in CL species was performed as described previously (Tatsuta, 2017) (1).

### dNTP pool measurement

dNTP pool measurement was done as previously described by (Nikkanen *et al*, 2016).

### Oxymetric analyses

Oxygen consumption rates were measured using a high-resolution oxygraph (OROBOROS instruments using a substrate-uncoupler-inhibitor protocol. Specific oxygen consumption rates are expressed as pmol/(s*mg). Briefly, 2×10^6^ cells were resuspended in respiration buffer (0.5 mM EGTA, 3 mM MgCl_2_, 60 mM Lactobionic acid, 20 mM Taurine, 10 mM KH_2_PO_4_, 20 mM HEPES, 110 mM D-Sucrose, 1% fat-free BSA). Oxygen consumption rates were performed in the presence of pyruvate-glutamate-malate and activities determined with +ADP (CI), +succinate (CI+CII), maximal uncoupled respiration by FCCP titration and CIV by ascorbate +TMPD.

### Statistical analysis

Statistical analysis data are present as the means ±SD, if not stated otherwise. Unless stated otherwise statistical significance was calculated using unpaired Statistical significance was determined using one–way ANOVA for mouse groups or Student’s t-test and represented in GraphPad Prism 8.4.1. In each figure sample size, repetitions and statistical test are justified. For false positive analysis of targeted metabolomics, the Benjamini-Hochberg method with a critical value of 0.2 was used.

## Author contribution

C.B.J., C.J.C., A.S. conceived and designed the study. C.B.J., A.M., T.M., F.S., L.E., N.Z., T.T., A.E.V., L.W., C.J.C. conducted the experiments. C.B.J., A.M., N.Z., T.T., A.E.V., L.W., R.A., analysed the data. B.J.B, T.L. guided and facilitated experiments. All authors edited the manuscript. C.B.J, A.M., A.S., wrote the manuscript, approved by all authors. A.S. supervised the study.

## Acknowledgments and Funding

The authors wish to thank Markus Innilä, Tuula Manninen and Babette Hollmann, for technical contributions and expertise; Prof. Johannes Spelbrink for the provision of the T-REX HEK POLG and TWNK overexpressor cell lines. Following funding resources are acknowledged: Academy of Finland [AS, CJ], Sigrid Jusélius Foundation [AS], Swiss National Science Foundation [CJ], the Magnus Ehrnrooth Foundation [CJ] and the Novartis Foundation for Medical-Biological research [CJ], Henry Wellcome Fellowship (215888/Z/19/Z) [AEV].

## The Paper Explained

### PROBLEM

To explore the role of serine biosynthesis as an integral component of stress response to mitochondrial translation disease and to evaluate it as a target for therapy.

### RESULTS

Despite serine intake via nutrition, we find serine to become an essential amino acid in mitochondrial stress. Pharmacological inhibition of *de novo* serine synthesis aggravates disease hallmarks, challenging particularly oxidative stress capacity and lipid synthesis affected.

### CLINICAL IMPACT

These results suggest that increasing *de novo* serine synthesis alleviates mitochondrial disease progression, with therapeutic implications.

## Supplementary figures

**Figure S1 (related to Figure 1).**
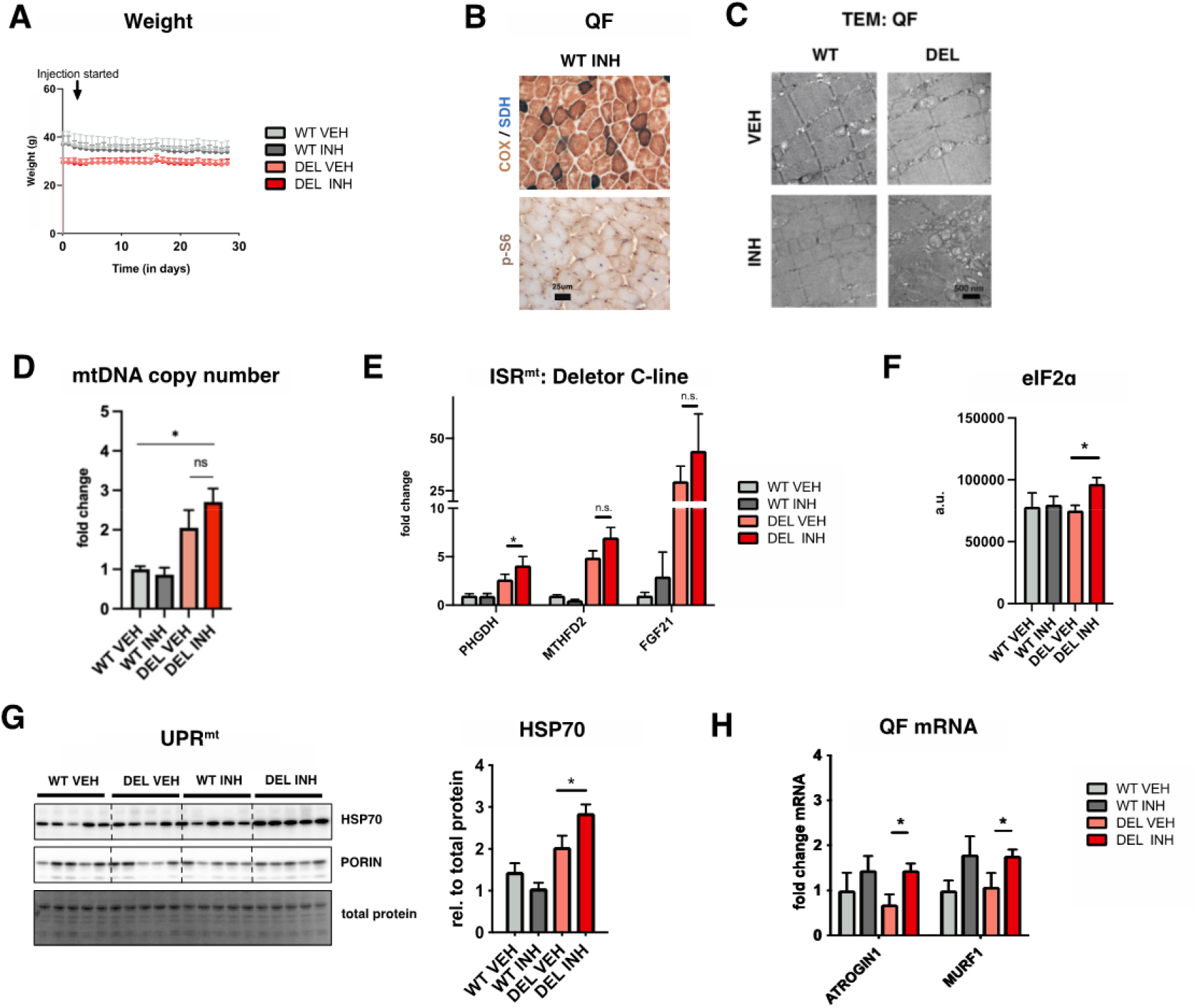
(A) Weight change over time of treatment with NCT-503 (WT VEH n=6, Del VEH n=5, WT INH n=6, Del INH n=8). (B) Histochemical analysis of combined cytochrome-c-oxidase (COX) and succinate dehydrogenase (SDH) activity in treated WT mice. (Brown fibres indicate high COX activity, translucent low COX activity. Lower panel shows immunohistochemical detection of the mTORC1 downstream target phosphorylated ribosomal S6 (S6-P). (C) Electron microscopic analysis. Left panel shows distorted mitochondria in vehicle and inhibitor-treated Deletors. (D) Mitochondrial DNA copy number analysis, qPCR (n=5-8/group). (E) Transcriptional upregulation of ISR^mt^ markers PHGDH, MTHFD2 and FGF21 in NCT-503 treated C-line Deletors. qPCR (n=4-6/group). (F) Quantification of immunoblot of phosphorylated eIF2α / total eIF2α (n=5/group). (G) Immunoblot of stress marker HSP70/GRP75 and mitochondrial mass marker with respective quantification for HSP70 relative porin (n=5/group). (H) Transcriptional levels of ATROGIN1, MURF1. qPCR (n=5-8/group).

**Figure S2 (related to Figure 2).**
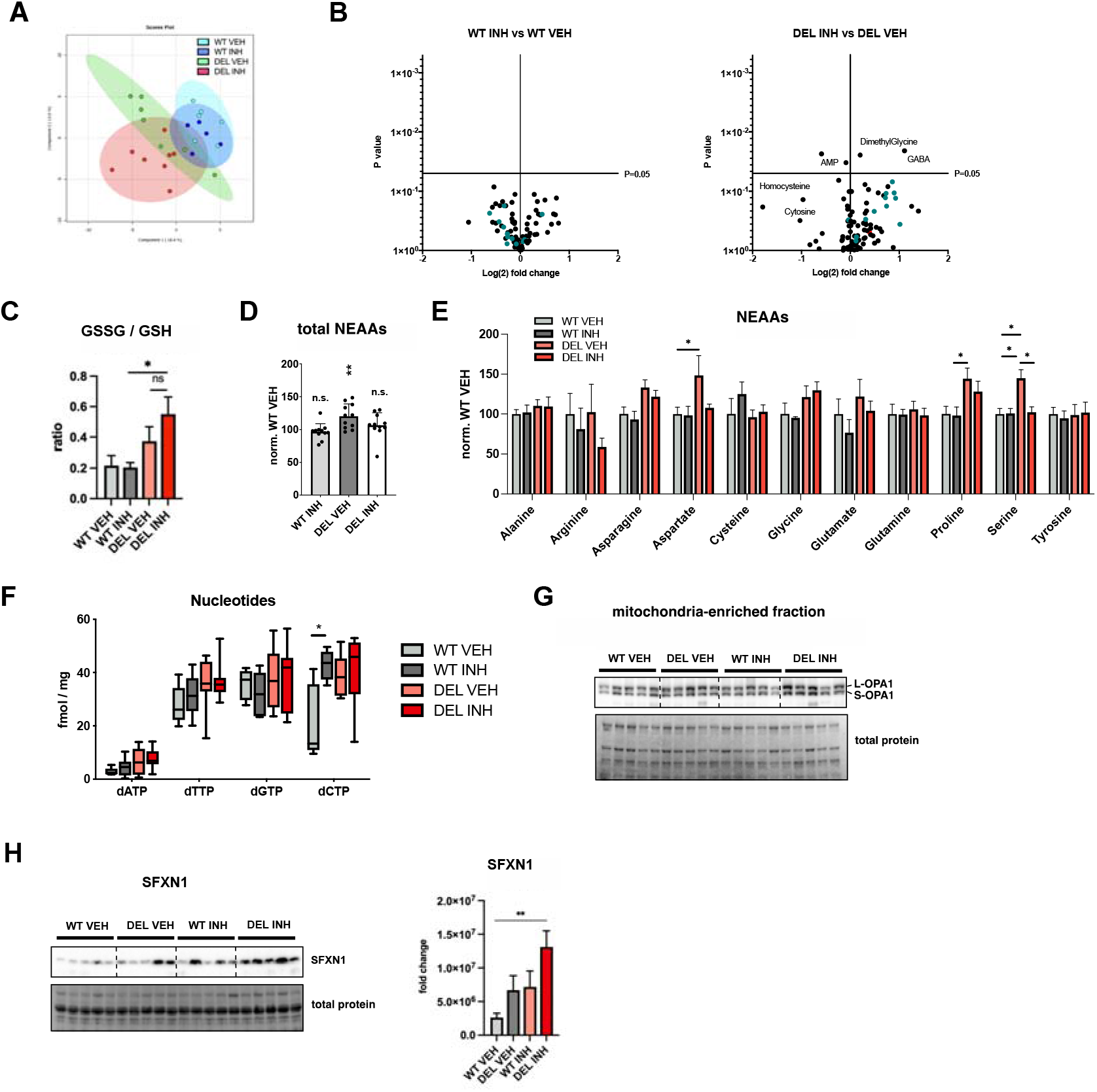
(A) PLSDA-plot of targeted metabolomics from skeletal muscle (WT VEH n=6, Del VEH n=5, WT INH n=7, Del INH n=8). (B) Volcano plots of targeted metabolomics from skeletal muscle. Dots represent individual metabolites. Acyl-carnitines are marked in green. Serine is marked in red. (WT VEH n=6, Del VEH n=5, WT INH n=7, Del INH n=8). (C) Ratio of oxidized (GSSG) and non-oxidised glutathione from (B). (D) Total non-essential amino acid quantification (NEAAs) from (B). (E) Individual non-essential amino acid quantification (NEAAs) from (B). (F) Total nucleotide pools in skeletal muscle. Total dNTP pools (WT VEH n=5, Del VEH n=7, WT INH n=6, Del INH n=8). (G) Immunoblot of isolated mitochondria from skeletal muscle for OPA1 (n=5/group). (H) Immunoblot of total muscle for SFXN1 with quantification (n=5/group).

**Figure S3 (related to Figure 3).**
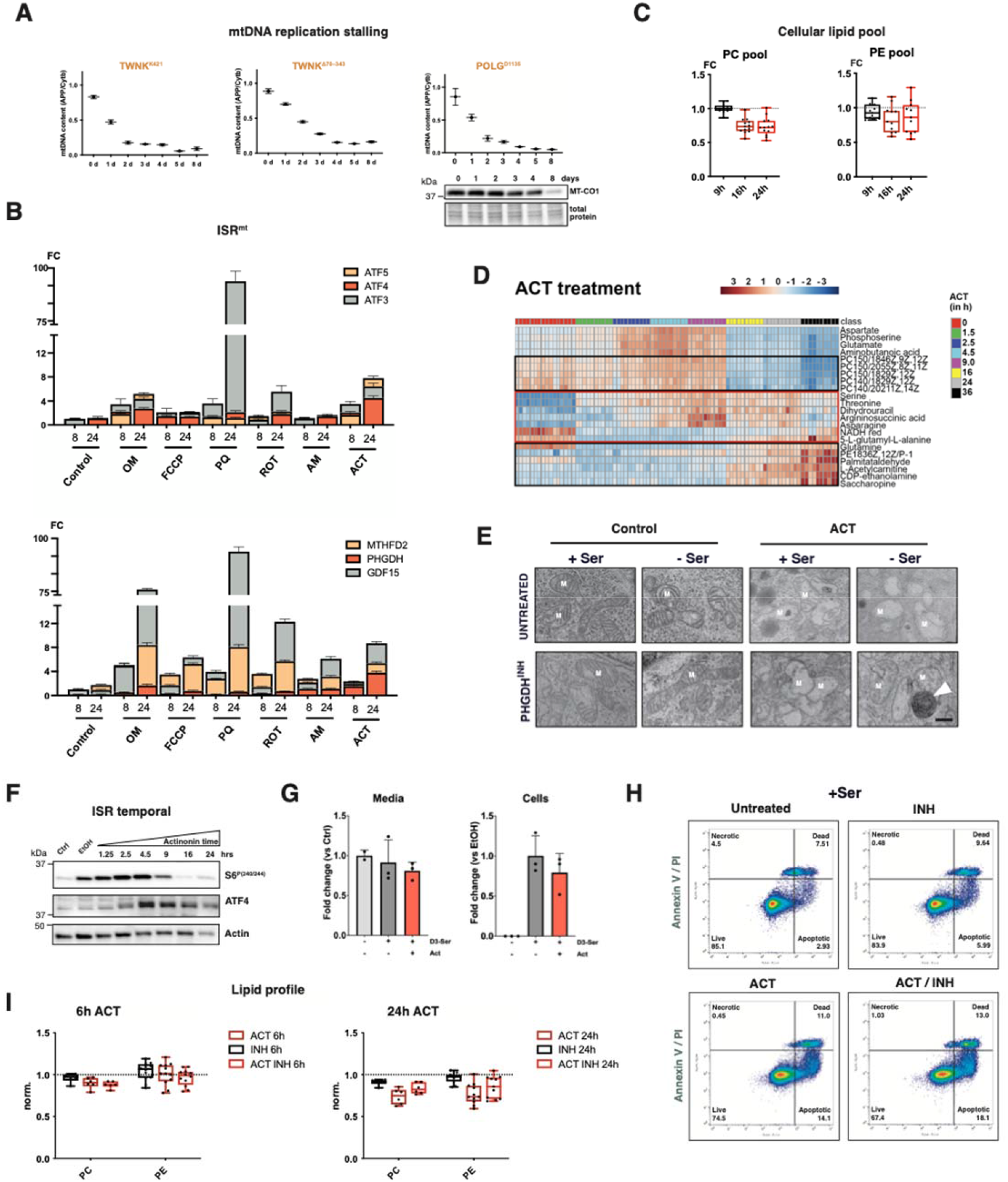
(A) MtDNA copy number dynamics upon overexpression of catalytic Twinkle (TWNK)- and DNA-polymerase gamma (POLG) mutants (TWNK^K421A^, TWNK^Δ70–343^, POLG^D1335A^ (Wanrooij *et al*, 2007)); qPCR analysis. HEK293T cells. POLG^D1135A^ mutant overexpression in HEK293T cells shows gradual decrease in mtDNA copy number with concomitant but lagging loss of MT-CO1. (B) Transcript levels of associated ISR^mt^ genes, qPCR (C) Total lipid pools including PC and PE species actinonin-treated C2C12 cells for the indicated time; untargeted temporal metabolomics. Each dot represents the median of a lipid species. (D) Heatmap of selected top significantly altered metabolites; untargeted temporal metabolomics. (E) Electron microscopic analysis of 12h treated cells. (F) Immunoblot of temporal actinonin-treatment for ATF4 and mTOR target phospho-S6 (p-S6). (G) D^3^-serine uptake. Amount in media and in cells; biological replicates (n=3). (H) Cell viability in serine-containing media. FACS (Annexin V / PI) Ctrl: live (85.5%) / apoptotic (2.58%) / dead (7.51%) INH: live (83.9%) / apoptotic (5.99%) / dead (9.64%) ACT: live (74.5%) / apoptotic (14.1%) / dead (11%) ACT/INH: live (67.4%) / apoptotic (18.4%) / dead (13.1%) (I) Normalised PC/PE pools, untargeted metabolomics. Abbreviations: PHGDH=D-3-phosphoglycerate dehydrogenase; ACT= actinonin, INH= NCT-503.

**Figure S4 (related to Figure 3).**
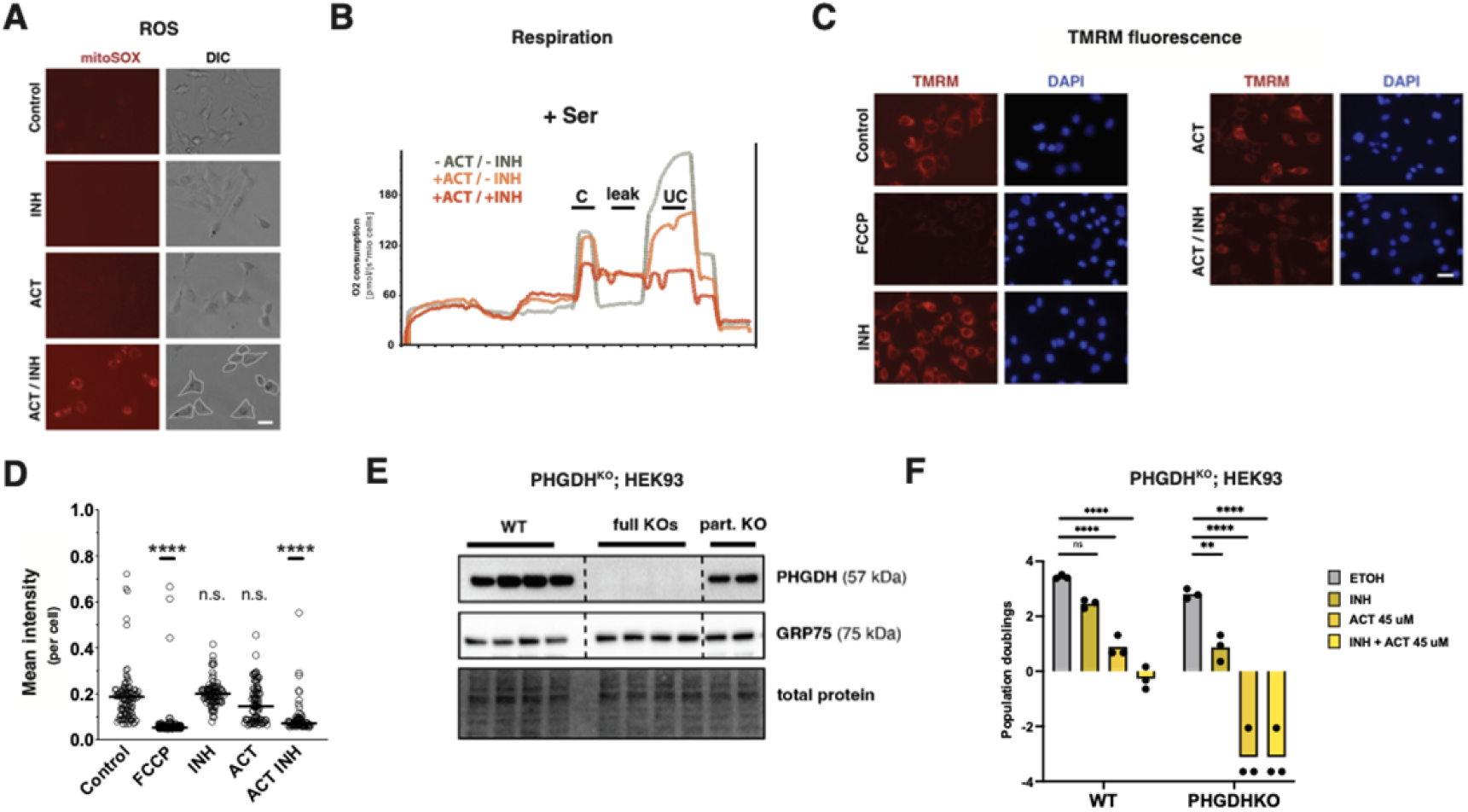
(A) Mitochondrial ROS levels upon inhibition of mt-translation and/or *de novo* serine biosynthesis. mitoSOX-probe detection (cell borders indicated with white dotted line in ACT /INH panel only). Scale bar: 20uM. (B) Respiration traces in permeabilised cells grown in actinonin and/or under PHGDH inhibition. C = coupled, UC = uncoupled. (C) Mitochondrial membrane potential upon inhibition of mt-translation and/or *de novo* serine biosynthesis. TMRM-probe detection, DAPI for cell count. Scale bar: 40uM. (D) Quantification of (C). (E) Immunoblot confirmation of PHGDH knockout(^KO^) in HEK293 cells. (F) Population doublings of WT and PHGDH^KO^ cells, three independent experiments.

**Table S1.**
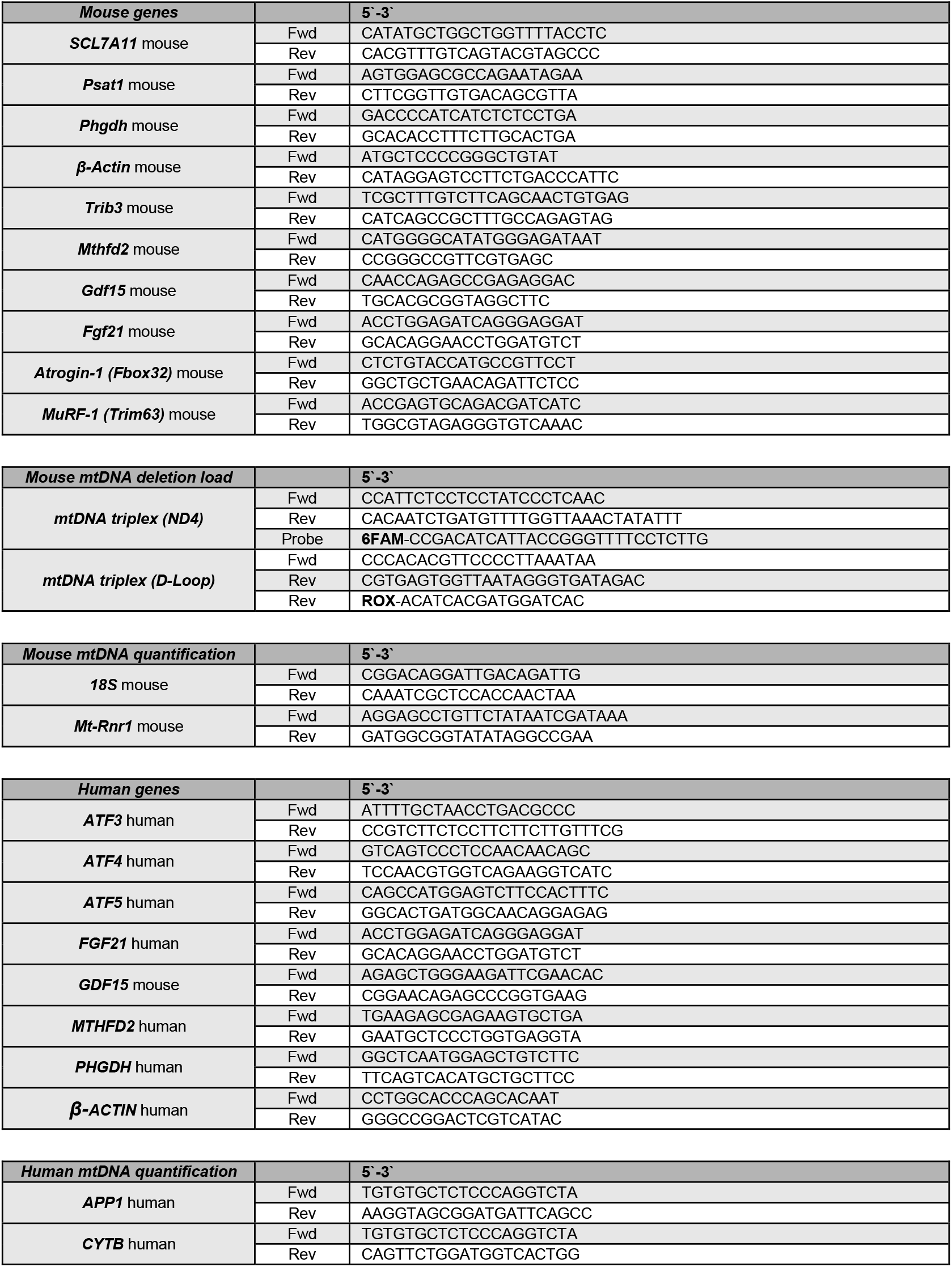
Primer sequences.

